# Oligogalacturonides, produced by enzymatic degradation of sugar beet by-products, confer partial protection against wheat powdery mildew

**DOI:** 10.1101/2024.11.15.623731

**Authors:** Camille Carton, Josip Šafran, Sangeetha Mohanaraj, Romain Roulard, Jean-Marc Domon, Solène Bassard, Natacha Facon, Benoît Tisserant, Gaelle Mongelard, Laurent Gutierrez, Béatrice Randoux, Maryline Magnin-Robert, Jérôme Pelloux, Corinne Pau-Roblot, Anissa Lounès-Hadj Sahraoui

## Abstract

Nowadays, it becomes urgent to use new alternative strategies for a more sustainable agriculture, by developing eco-friendly biocontrol compounds leading to an induction of plant resistance against pathogens. In recent years, oligogalacturonides (OGs), fragments of pectin resulting from the degradation of the plant cell wall, have been particularly under focus for their ability to induce plant defense, leading to plant protection against pathogens. In the current work, 2 mixtures of OGs, bio-sourced from sugar beet by-products, were produced and assessed for their protective efficacy on wheat against *Blumeria graminis f. sp. tritici* (*Bgt*), responsible for powdery mildew. These OGs pools were obtained by hydrolysing pectins with two *A. thaliana* polygalacturonases (PGs), ADPG2 and PGLR (EC 3.2.1.15), produced in heterologous system *Pichia pastoris*. No direct effect of OGs generated by either ADPG2, named OGs-ADPG2 (degree of polymerisation/DP 1-10 and degree of esterification/DE 41%) or PGLR, named OGs-PGLR (DP 1-16, DE 34 %), was observed on *Bgt* spore germination *in vitro*. After a preventive foliar spraying of wheat by OGs, only OGs-PGLR provided a 31 % partial protection against *Bgt*. Both pools of OGs triggered an increase in the expression of some plant defense-related genes (peroxidase or PR proteins encoding genes), before inoculation with the pathogen. However, only OGs-PGLR induced the expression of a gene encoding a lipoxygenase (LOX), which is the first enzyme involved in the octadecanoic pathway, leading to oxylipins production in plant defense. Taking together, these findings revealed that some OGs, produced from sugar beet by-products by enzymatic hydrolysis, could be used as an alternative of phytochemicals.

## 1. Introduction

Bread wheat (*Triticum aestivum L*.) is one of the most predominant cereal crops in the world (Kang et al., 2020; Ochoa-Meza et al., 2021). It plays an important role in global food security since the Green Revolution (Curtis and Halford, 2014; Ochoa-Meza et al., 2021). In 2020, wheat harvest was approximately 757.6 million tons (Kang et al., 2020; Kumar et al., 2024; Ochoa-Meza et al., 2021). However, wheat can be subjected to various diseases, caused by fungal and bacterial agents, leading to a loss in grain yield and quality (Mwadzingeni et al., 2016).

*Blumeria graminis f. sp. tritici* (*Bgt*) is an obligate biotrophic pathogenic fungus, responsible for powdery mildew in wheat, that causes significant damages leading to loss of almost 5 % of the global annual wheat yield (Zou et al., 2023). Once this disease is established in a wheat-growing area, production losses can reach up to 35, 40 and 62 % in Russia, China, and Brazil, respectively (Kang et al., 2020; Zou et al., 2023). Currently, the use of fungicides and the selection of more resistant cultivars are effective strategies for managing this disease (Bartlett et al., 2002; Vielba-Fernández et al., 2020). However, the use of chemicals is increasingly controversial as they are potentially harmful both to human health and to the environment. Moreover, the widespread use of fungicides has resulted in the development of resistances in various fungal populations, leading to a reduction in their efficiency (Godet and Limpert, 1998; Meyers et al., 2019). Therefore, to promote a more sustainable agriculture and safer food production, it is important to develop new eco-friendly disease-control alternatives. Thus, the development of bio-based resistance inducers, substances that trigger plant defense mechanisms against pathogens, plays a crucial role in enhancing crop resilience and reducing the need for chemical pesticides. Plant resistance inducers, called also elicitors, are one of the most promising alternatives that meet these requirements (Thakur and Sohal, 2013).

Cell wall represents one of the first plant defense barrier against pathogens. Despite the vast diversity of plant species, their primary cell wall shares a similar composition. This extracellular layer is constituted of 3 distinctive polysaccharides: cellulose, hemicellulose and pectins (Zhang et al., 2021). During pathogen infection, plant cell wall may serve as extracellular sources of damage-associated molecular patterns (DAMPs) by releasing polysaccharides fragments (Pontiggia et al., 2020). DAMPs carbohydrates include cellulose-derived cellobiose and cellotriose, mixed-linkage glucans, and pectin-derived oligogalacturonides (OGs) (Bacete et al., 2018). Pectins are composed of four heterogeneous polysaccharides: xylogalacturonan (XGA), rhamnogalacturonan II (RG-II), rhamnogalacturonan I (RG-I) and homogalacturonan (HG) which is generally the most abundant (Shahin et al., 2023). HG is an homopolymer of α-1,4-linked-D-galacturonic acid (GalA) units, where methylester and/or acetyl groups may be present (Mohnen, 2008). The depolymerization of HGs in short chains of GalA residues produce OGs, which are natural plant elicitors (Ferrari et al., 2013). OGs are classified with an elicitor activity (Albersheim et al., 1983), and are naturally produced during infection by a pathogen (Ferrari et al., 2013). OGs are able to activate signalling pathways mediated by Ca^2+^ leading to the activation of plant defense responses such as deposition of callose (Galletti et al., 2008; Gravino et al., 2015), production of reactive oxygen species (ROS) (Gravino et al., 2015; Lecourieux et al., 2002; Navazio et al., 2002) and nitric oxide (Rasul et al., 2012).

OGs can be produced by enzymes such as polygalacturonases (PGs) present in plants and pathogenic fungi, bacteria or nematodes (Benedetti et al., 2017; Ferrari et al., 2013; Silva-Sanzana et al., 2020). The activity of these enzymes depends on the structure of the HG including it degree esterification (DE), such as degree of methylation (DM) and degree of acetylation (DA). Thus, the hydrolysis of HGs by PGs can produce of OGs of peculiar structures, including their degrees of polymerization (DP) and esterification. However, few studies exploited the natural esterification of substrates and none used bio-based substrates to produce OGs. Thus, being able to produce esterified OGs, in a controlled manner and without thermal of chemical treatment, is one of the objectives of more sustainable agriculture.

The exogenous application of OGs on plants, under controlled conditions, triggered resistance against a wide range of pathogenic fungi in various plant species: (i) OGs with a DP of 2–7 and an undetermined esterification level conferred resistance to *Rhizoctonia solani* in sugar beet roots (Zhao et al., 2022); (ii) non-esterified OGs with a DP of 10 to 15 protected tomato plants against *Botrytis cinerea* (Gamir et al., 2021); and (iii) OGs with a DP of 2 to 25, chemically acetylated at 30%, were also shown to protect wheat from powdery mildew (Randoux et al., 2010). Additionally, the application of OGs against *Bgt* demonstrated the induction of enzymatic activities such as oxalate oxidase, peroxidase, and lipoxygenase, which are involved in the establishment of plant defences (Randoux et al., 2010).

The use of pectin-rich agricultural by-products, such as citrus peels (Willats et al., 2006), apple pomace (Putra et al., 2023), grape pomace (Spinei and Oroian, 2023, 2022) and sugar beet pulp (Joanna et al., 2018), has not been explored until now for OGs production. After sucrose extraction, approximately 80% of the sugar beet weight is converted into pulp, which is a resource rich in pectin. This pectin, naturally methylated and acetylated (Zeuner et al., 2020), could be interesting for the production of OGs that could be tested for plant protection against phytopathogens, such as *Bgt* on wheat.

Our study therefore aims to produce OGs from sugar beet pulp by enzymatic degradation with PGs, while maintaining the characteristics of this bio-based substrate (DM and DA). Moreover, our method aims to reduce thermal and chemical treatments to align with a sustainable process. First, the structure of the pectins were determined by nuclear magnetic resonance (NMR) spectroscopy and high-performance anion exchange chromatography (HPAEC). Using two PGs from *Arabidopsis thaliana*, ADPG2 and PGLR, produced in heterologous system *Pichia pastoris,* two pools of OGs were obtained by enzyme degradation of sugar beet pectins (SBP), with various DP, DM and DA. The OGs-ADPG2 and OGs-PGLR were further tested as a preventive treatment against wheat powdery mildew. To understand the mode of action of both pools of OGs, their potential direct antifungal activity was first assessed by using *in vitro* antigerminative tests on *Bgt* spores. Their potential elicitor activity was studied by measuring the expression of defense-related genes in wheat leaves after preventive treatments with both pools of OGs and inoculation with *Bgt*.

## 2. Materials and Methods

### 2.1. Production and determination of enzymatic activity of polygalacturonases

#### 2.1.1. Cloning and heterologous expression of ADPG2 and PGLR

*A. thaliana* polygalacturonases (EC 3.2.1.15) ADPG2 and PGLR were previously cloned and expressed in the yeast *P. pastoris* (Hocq et al., 2020; Safran et al., 2023).

#### 2.1.2. Production and purification of the recombinant ADPG2 and PGLR proteins

Yeast cells were incubated at 30 °C and 250 rpm in baffled flasks containing buffered glycerol-complex medium (1 % yeast extract, 2 % peptone, 100 mM potassium phosphate pH 6, 1.34 % YNB, 4×10^-5^ % biotin and 1 % glycerol). After 24 hours, the cells were transferred to buffered methanol-complex medium without glycerol, with a final methanol concentration of 0.5 %. Methanol concentration was maintained by adding 2 mL of 25 % methanol to 100 mL of culture every 24 hours, ensuring a constant concentration of 0.5 % methanol in the medium over 72 hours at 30°C and 250 rpm (Lemaire et al., 2020). The culture was then centrifuged (1,500 g, 10 min, 4 °C) and filtered using GD/X 0.45 μm PES filter Media (Whatman, Maidstone, United Kingdom).

Purification of ADPG2 and PGLR proteins was conducted using a 1 mL HisTrap excel column (GE Healthcare, Chicago, Illinois, United States). The column was initially equilibrated with 10 volumes of equilibration buffer (50 mM NaP pH 7.2, 250 mM NaCl). Next, 100 mL of supernatant culture was loaded into the column at a flow rate of 1 mL.min^-1^. The HisTrap column was washed with 10 column volumes of wash buffer (50 mM NaP pH 7.2, 250 mM NaCl, 25 mM imidazole). The recombinant protein was eluted by the addition of 10 column volumes of elution buffer (50 mM NaP pH 7.2, 250 mM NaCl, 500 mM imidazole). ADPG2 and PGLR proteins were concentrated using an Amicon Ultra Centrifugal filter with a 10 kDa cut-off (Merck Millipore, Burlington, Massachusetts, United States). Buffer exchange of ADPG2 and PGLR proteins was performed using a PD Spintrap G-25 column (GE Healthcare, Tremblay-en-France, France) in ammonium acetate 50 mM pH 5.2.

#### 2.1.3. Determination of PG concentration and activity

Protein concentrations were assessed using the Bradford assay (Bradford, 1976), with Bovine Serum Albumin (A7906, Sigma-Aldrich, Saint-Quentin-Fallavier, France) as a standard. Subsequently, PGs were diluted to a concentration of 1 µg.µL^-1^, and the activities of purified ADPG2 and PGLR enzymes were determined using the DNS method (Miller, 1959) in 50 mM ammonium acetate buffer at pH 5. The reaction mixture contained polygalacturonic acid (PGA) (81325, Sigma-Aldrich, Saint-Quentin-Fallavier, France) at a final concentration of 0.4 % and was incubated at 50 °C for 60 minutes.

### 2.2. Production and characterisation of Oligogalacturonides

#### 2.2.1. Production and characterisation of pectic substrate

SBP was obtained from beet pulp recovered after extraction of sucrose and finely ground using a ball mill. Pectins was isolated by treating the pulp with 50 mM HCl adjusted to pH 5 (with NH_4_OH 1M) in a water bath for 1 h at 70 °C. After centrifugation 15 min at 2500 rpm, the supernatant was preserved and the pectins were precipitated by adding 3 volumes of 96 % ethanol overnight at 4 °C. Then, the mixture was centrifuged for 15 min at 4 °C at 2500 rpm. The supernatant was eliminated, and the pectin pellet was solubilized in ultra-pure water, frozen then dried by freeze-drying.

The monosaccharide composition of pectins was determined after hydrolysis (1 mg.mL^-1^) with 4 M CF_3_COOH (100 °C for 4 h). Aliquots of the extract were analyzed by high-performance anion exchange chromatography (HPAEC) equipped with a pulsed amperometric detector (Dionex^TM^ ICS 5000+ system) by injecting 2.5 µL on 2 x 50 mm guard column and 2 x 250 mm column CarboPac™ PA1 at 30 °C with 0.25 mL.min^-1^ flow rate. A multi-step gradients elution was carried out as follows: for neutral monosaccharides: 0-25 min, sodium hydroxide (NaOH) 16 mM; 26-35 min, NaOH 200 mM; 36-50 min, NaOH 16 m (4mM replacing 16mM NaOH were used to separate xylose and mannose as they co-elute in this usual method with carboPac PA1), and for uronic acids: 0-5 min, 100 % NaOH 160mM; 5-35 min 0 to 100 % NaOH 160mM with sodium acetate 600mM maintained until 40 min, 42-55 min 100 % NaOH 160mM. Peak processing and analysis were performed using the Cobra Wizard algorithm (Chromeleon™ 7.2.10 software).

For nuclear magnetic resonance (NMR) spectroscopy, pectins were dissolved in 99.96 % D_2_O (10 mg.mL^-1^). ^1^H NMR spectra were recorded at 50 °C, in order to reduce the viscosity, on a Bruker Avance 500 spectrometer equipped with a BBI probe (5-mm sample diameter) and Topspin 1.3 software. ^1^H NMR spectra were accumulated using a 30° pulse angle, a recycle time of 1 s, and an acquisition time of 2 s for a spectral width of 3,000 Hz for 32-K data points using an experimental sequence provided by Bruker.

The degree of methylation (DM) and the degree of acetylation (DA) of SBP were determined by quantification of methanol and acetate after saponification of pectin extracts. Pectins were dissolved in D_2_O, and a first ^1^H NMR experiment at 50 °C was performed in order to check the absence of free methanol and acetate. Then, 100 µL of NaOD (1 M) in D_2_O were added in the NMR tube, and spectrum was further recorded 10 min after the addition of NaOD. Under the conditions applied, saponification was complete, and no methanol evaporation was observed. The DM and DA values were then calculated as described previously (Bédouet et al., 2003) and expressed in percent per residue.

#### 2.2.2. Production of oligogalacturonides (OGs)

Sugar beet pectin extract was dissolved in 50 mM ammonium acetate buffer pH 5.2 at a concentration of 0.8 % (w/v). Purified ADPG2 or PGLR were diluted to reach total activities of 1.2 nmol.min^-1^.μg^-1^ for ADPG2 and 0.8 nmol.min^-1^.μg^-1^ for PGLR, and 10 % (v/v) of these enzyme solutions were added to substrates for hydrolysis. Pectin hydrolysis was carried out at a concentration of 0.4% (m/v) for 2 h in 50 mM ammonium acetate buffer pH 5.2. The reaction was stopped by adding one volume (v/v) of absolute ethanol and the mixtures were kept overnight with stirring at 4 °C. The supernatant was collected after centrifugation at 4 °C for 5 min at 5000 g and dried using a SpeedVaq (Eppendorf, Montesson, France). Then, the OGs were solubilized in LC/MS water, frozen and dried again by freeze-drying to obtain a powder. The OGs produced by hydrolysis of SPB by ADPG2 were noted: OGs-ADPG2 and those produced by hydrolysis of sugar SPB by PGLR were noted: OGs-PGLR.

#### 2.2.3. OGs characterization

First, the OGs produced as described in section 2.2.2 were characterized by ^1^H NMR at room temperature (2.2.1). Then, the OGs were suspended in 200 µL in LC/MS water at 5 g.L^-1^. Chromatographic separations of OGs were performed using an ACQUITY UPLC Protein BEH SEC column (125Å, 1.7 μm, 4.6 mm x 300 mm) and identification of OGs were performed by high resolution electrospray mass spectrometry (ESI-HRMS) with the SYNAPT G2-Si instrument coupled to the ACQUITY UPLC H-Class system with the method described by Voxeur et al. (2019).

### 2.3. Application of oligogalacturonides on wheat

2.3.1. Plant treatments, and inoculation with *Blumeria graminis f. sp. tritici* (*Bgt)*

The experiments were carried out on bread wheat, *Triticum aestivum*, cultivar Alixan (Limagrain, Saint-Beauzire, France). This cultivar presents a resistance level against powdery mildew estimated in field at 6, on a resistance scale from 1, for fully susceptible and 9, for fully resistant cultivars. The grains were sowed in compost (Floragard, Oldenburg, Germany) in trays (24 grains per tray) and then transferred to a growth chamber, with a 12 h photoperiod (with a light intensity of 250 µmol.m^−2^.s^−1^), a relative humidity of 70 %, day temperatures of 18 °C and night temperatures of 12 °C. Treatment with OGs or water-control occurred when plantlets were 16 days old (**Fig. S1**). OGs-PGLR and OGs-ADPG2 powders were dissolved at 5 g.L^-1^ in 10 mL of dH_2_O and sprayed on wheat plantlets as described by Randoux et al. (2010). The wheat leaves were inoculated with 8 days-old conidia of *Bgt*, collected from fully infected leaves by vacuum extraction, 2 days after treatment with OGs. These conidia were suspended in Fluorinert FC43 (3M, Cergy-Pontoise, France), a product compatible with the hydrophobic nature of spores and having no impact on either the plant or the fungus. The inoculum concentration was adjusted to 5×10^5^ spores.mL^-1^, and 7.5 mL of the conidia solution were sprayed onto the leaves of 24 plants.

#### 2.3.2. Symptoms evaluation

Nine days after inoculation with *Bgt* conidia, 14 plantlets, whether treated with OGs or not, were harvested to count the number of colonies that had developed on the adaxial and abaxial surfaces of the third leaf. The percentage of infection achieved with OGs solutions was calculated using the following formula: (average number of colonies on treated leaves ÷ average number of colonies on control leaves) × 100.

#### 2.3.3. *In vitro* antifungal assay

Following a first screening, OGs-ADPG2 and OGs-PGLR were tested at 5 g.L^-1^ diluted in dH_2_O as described by Randoux et al. (2010). Petri dishes were filled with agar at 12 g.L^-1^, prepared in dH_2_O, and 200 µL of OGs solution were spread on the medium, the control being an agar medium containing only deionized water. Fresh spores of *Bgt* were dispersed on the dishes and then observed after 24 h of incubation at 20°C. The germination was randomly determined on 200 conidia per plate using a light microscope (Olympus BX 40). The conidia were classified into five categories: non-germinated spores, spores with aborted germ tubes, spores with a short germ tube, spores with a long germ tube, and spores with multiple germ tubes. The distribution of spores across these five categories was reported by their frequency.

#### 2.3.4. RNA extraction, cDNA synthesis and RT-qPCR gene expression analysis

In order to assess the modifications in expression levels of defense genes induced by the different treatments (**Fig. S1**), wheat leaf samples were harvested 48 h post-treatment (hpt) with OGs or dH_2_O (before *Bgt* inoculation), as well as 24 h (D1) and 48 h (D2) post-inoculation with *Bgt* (equivalent to 72 and 96 hpt under non-infectious conditions). For each condition, the third leaf from three wheat plantlets was harvested, flash-frozen in liquid nitrogen and ground into powder. Total RNA was extracted from 100 mg of plant tissue using the RNeasy Plant Mini Kit (Qiagen, Hilden, Germany) according to the manufacturer’s instructions. The resulting RNA preparations were treated with DNaseI using the RNase-Free DNase Set (Qiagen, Hilden, Germany).

Two μg of RNA were reverse-transcribed using the High Capacity cDNA Reverse Transcription Kit (Applied Biosystems, Waltham, USA) according to the manufacturer’s protocol. Transcript levels were assessed in three independent biological replicates by real-time RT-qPCR with duplicate reaction mixtures containing 0.5 µM of both forward and reverse primers, cDNA, PowerUp™ SYBR® Green (Applied Biosystems Waltham, USA) and water. RT-qPCR was performed using QuantStudio™ 7 Flex Real-Time PCR System (Thermo Fisher Scientific, Waltham, USA).

Primers listed in **Table S1** were used to quantify the expression of 11 target genes (i.e. *chitinase – CHI, basic chitinase 1 – CHI1, acidic chitinase (class 4) – CHI4, Pathogenesis related protein 1 – PR1, Glucanase – GLU, Phenyalanine ammonia lyase – PAL, Peroxidase 2 – POX2, Peroxidase 381 – POX381, Glutathion - S - Transferase class phi – GSTphi, phosphoinositide specific Phospholipase – PIPCL2, Lipoxygenase – LOX*), which were selected from previous studies performed on wheat–pathogenic fungus interactions (Khong et al., 2013; Mustafa et al., 2017; Ors et al., 2018; Tayeh et al., 2014).

Normalization of RT-qPCR was performed using reference genes, which were selected and validated as follows. *TUB (Beta-tubulin 4)*, *ACT (Actin)*, *ATPAAA (ATPAse family AAA)* and *PETA (Apetala 2 class A)* genes were chosen for their putative stability of expression according to Tayeh et al. (2014) and; Velho et al. (2022). Their expression was assessed under our experimental conditions and these genes were ranked according to their stability of expression using geNorm software (Vandesompele et al., 2002). *TUB* and *ATPAAA* were the most stably expressed genes and thus were used to normalize the real-time RT-PCR data. The normalized expression patterns obtained using both reference genes were similar, so only the data normalized with *TUB* are shown in this article. Data analysis was performed using QuantStudio™ Design and Analysis v2. The crossing threshold values (CT) were used to calculate relative gene expression as in Lakehal et al. (2019). In brief, for each kinetic point, the gene expression values are relative to the expression in the respective control (untreated and uninfected), for which the value was set to 1.

#### 2.3.5. Statistical analysis

For the protection assay, a T-test with Welch’s correction was carried out in comparison with the control untreated plants. All the statistical analysis was carried out with the software GraphPad Prims version 8.

## 3. Results

### 3.1. Production and characterization of oligogalacturonides from pectin of sugar beet

Sugar beet production and the various industrial treatments of its pulp generate significant waste, which can be regarded as a valuable by-product containing cellulose, hemicellulose and pectin (Usmani et al., 2022). In this study, sugar beet pulp, after sucrose extraction, was utilized to extract pectins, which were then used as substrates for polygalacturonases (PGs) to generate oligogalacturonides (OGs)

The analysis of sugar beet pectin (SBP) composition by HPAEC showed a high content of neutral sugars, with arabinose (48 %), galactose (20%), rhamnose (6%) and glucose (5%). The acidic sugars represented 18% of total sugars and were mainly composed of galacturonic acid (17%) (**Fig. 1**). Similar proportions of arabinose, galactose and galacturonic acid were reported by Sakamoto and Sakai (1995). Additionally, the ^1^H NMR analysis (**Fig. 2A**) revealed characteristic spectra of pectins, with the presence of H-1 region between 4.8 and 5.2 ppm, signals of over proton of monosaccharides between 3 and 4.6 ppm and CH_3_ signal of C-6 of rhamnose residues at 1.30 ppm, characteristic of rhamnogalacturonan-I domain (RGI) (Winning et al., 2007). Several acetylesterified group signals were observed around 2 ppm indicating the presence of *O*-2 and/or *O*-3 acetylation on galacturonic acid residues, and an important signal of methylesterified group at 3.72 ppm was observed on spectra. After saponification, pectin was estimated to have 62% methyl esterification and 27% acetyl esterification. These results corroborate the composition of SBP previously detailed by Zeuner et al., (2020), highlighting its relatively high degree of methylesterification and acetylesterification. Therefore, SBP used in the current study seems to be an interesting candidate as PGs could generate bigger diversity of OGs (Ralet et al., 2005).

**Fig. 1.**
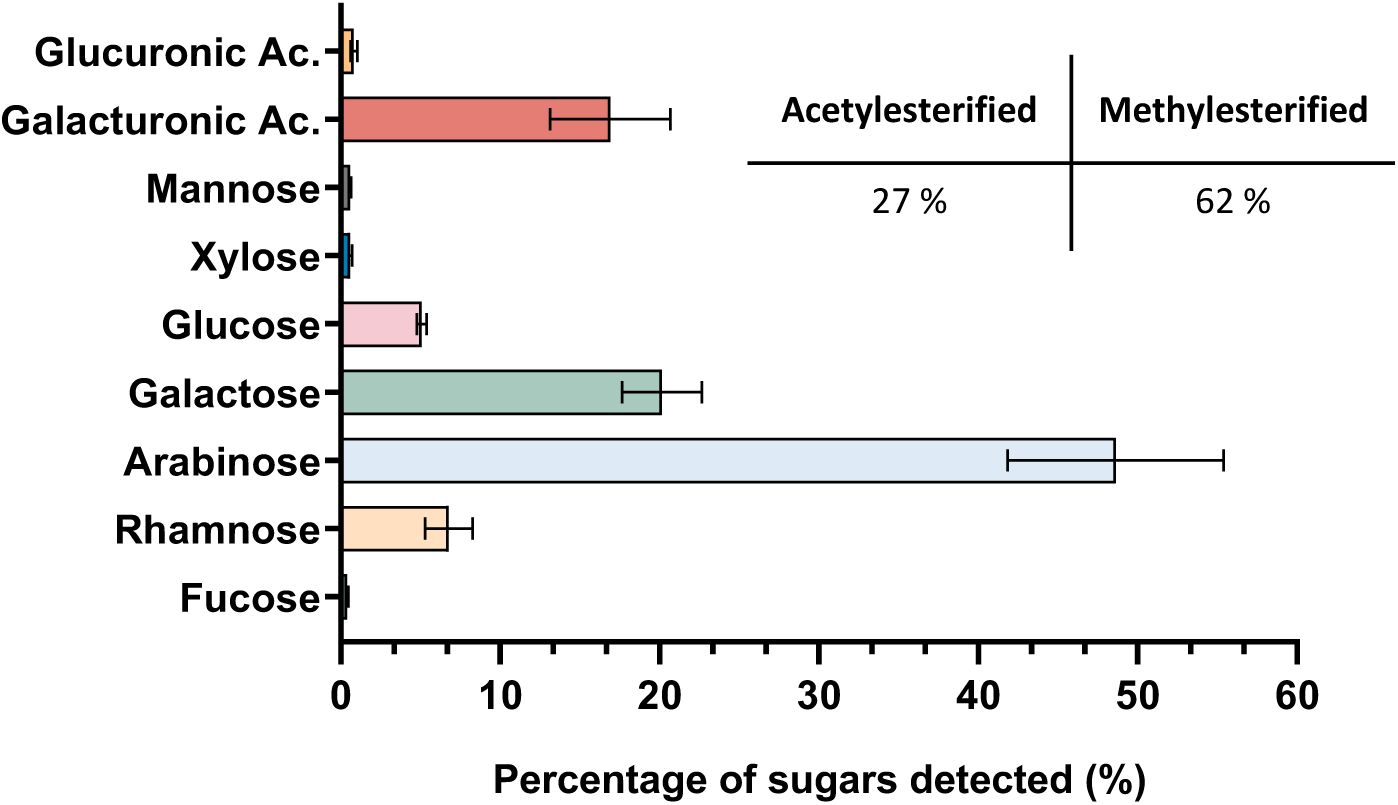
Neutral and acidic monosaccharides composition of the sugar beet pectin. Inset: Degree of esterification.

**Fig. 2.**
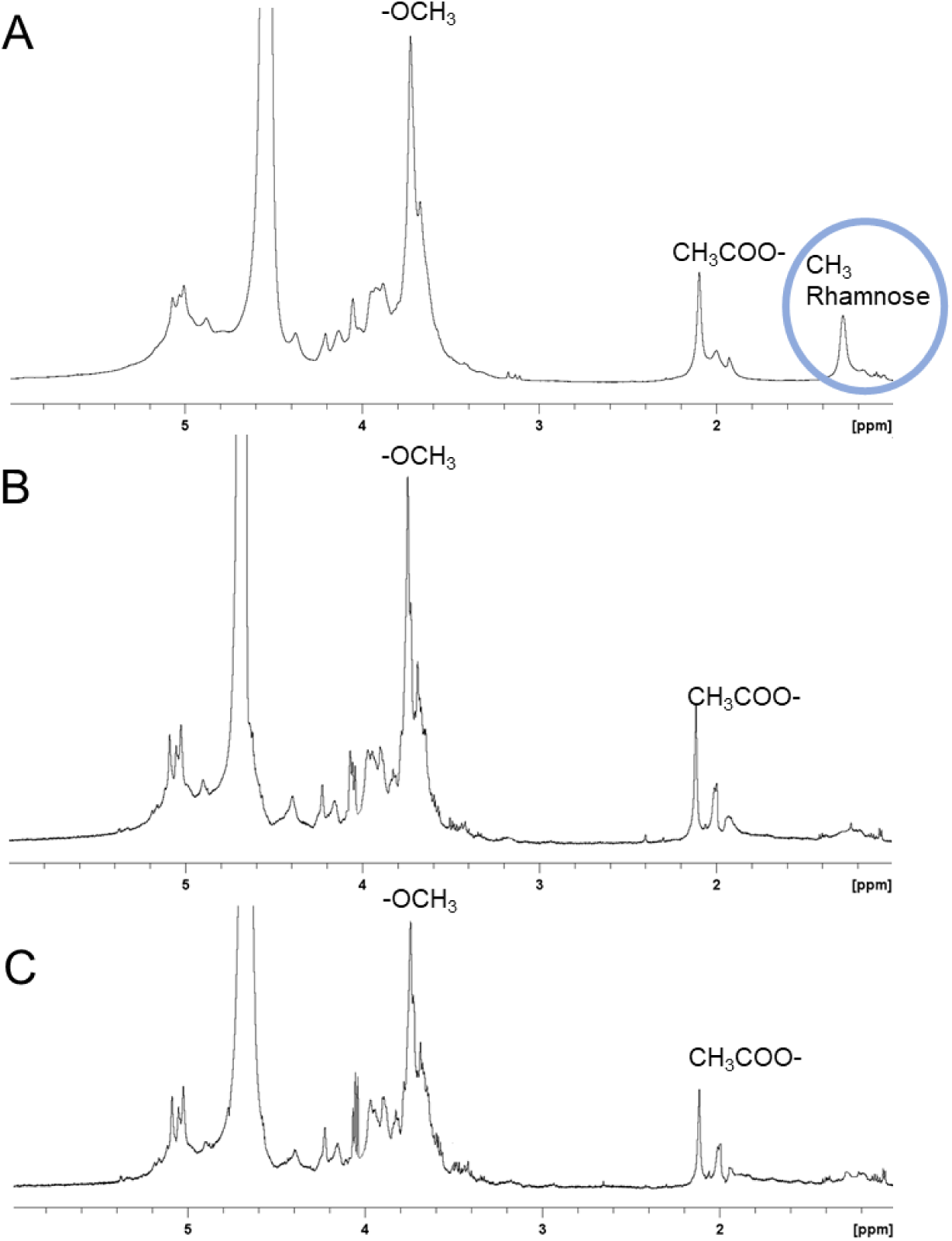
^1^H NMR spectra of sugar beet pectin (A) and oligogalacturonides production by ADPG2 (B) and PGLR (C).

OGs production was carried out with two endo-polygalacturonases, ADPG2 and PGLR from *A. thaliana.* The ability of these two endo-PGs to produce various OGs from HGs has already been demonstrated by Safran et al. (2023). Based on these results, we confirmed the structure of the OGs produced by ADPG2 and PGLR through ^1^H NMR analysis **(Fig. 2B and C**). Spectra showed a signal for methylesterified group at 3.72 ppm and also acetylesterified group signals around 2 ppm indicating that the OGs produced by PGs were esterified. The disappearance of the signal of CH_3_ of rhamnose at 1.3 ppm indicates that these residues were not present in OGs highlighting that RGI domains was not degraded by PGs and was eliminated during ethanol precipitation. These results confirmed that the conditions used allowed the production of OGs from SBP thanks to the hydrolysis of HG-type pectins by PGs. Moreover, methyl and acetyl ester group were preserved.

LC/MS oligoprofiling approach was used to identify the different OGs released during the 2 h hydrolysis of SBP by the two PGs (Voxeur et al., 2019). The results showed differences in OG pools following hydrolysis by PGLR and ADPG2 (**Fig. 3**). OGs produced had a DP between DP 1 to 10 for ADPG2 and PGLR while GalA16Me5 was additionally produced by PGLR (**Fig. 3A**). In both cases the OGs with the highest relative proportion is GalA3 (OGs with DP 3). Both OGs pools displayed various DP, with ADPG2 having a higher relative proportion of OGs of DP<4 (68%) as compared to PGLR (58%) (**Fig. 3B**). Indeed, we found that only 32 % of OGs-ADPG2 had a DP > 4, whereas 42 % of OGs-PGLR had a DP > 4 (**Fig. 3B**). These results were consistent with the characterization of Safran et al. (2023) showing that the OGs resulting from the hydrolysis by PGLR had a higher DP than those hydrolysed by ADPG2, indicating a difference in the mode of action of the enzymes. The OGs-ADPG2 had an overall degree of esterification of 41 %, which included 14 % methyl/acetyl groups, 1 % acetyl groups, and 26 % methyl groups (**Fig. 3C**). For OGs-PGLR, the overall de gree of esterification was 34.5 %, comprising 12.5 % methyl and acetyl groups, and 22 % methyl groups (**Fig. 3C**). Finally, we observed that certain OGs were specifically produced by either ADPG2 or PGLR (**Table S2**). For instance, ADPG2 specifically produced 15 OGs: GalA3Ac1, GalA4, GalA4Me2, GalA5Me1Ac1, GalA5Me2, GalA5Me3, GalA6Me3, GalA7Me4, GalA7Me4Ac1, GalA8Me4, GalA8Me4Ac2, GalA9Me6, GalA9Me6Ac1, GalA9Me6Ac2, and GalA10Me6. In contrast, PGLR specifically produced 16 OGs: GalA4, GalA5, GalA5Me1, GalA5Me2, GalA6, GalA6Me1, GalA7Me2, GalA7Me2Ac1, GalA7Me3, GalA7Me3Ac1, GalA9, GalA9Me4Ac1, GalA10, GalA10Me6Ac2, and GalA16Me5 (**Table S2**).

**Fig. 3.**
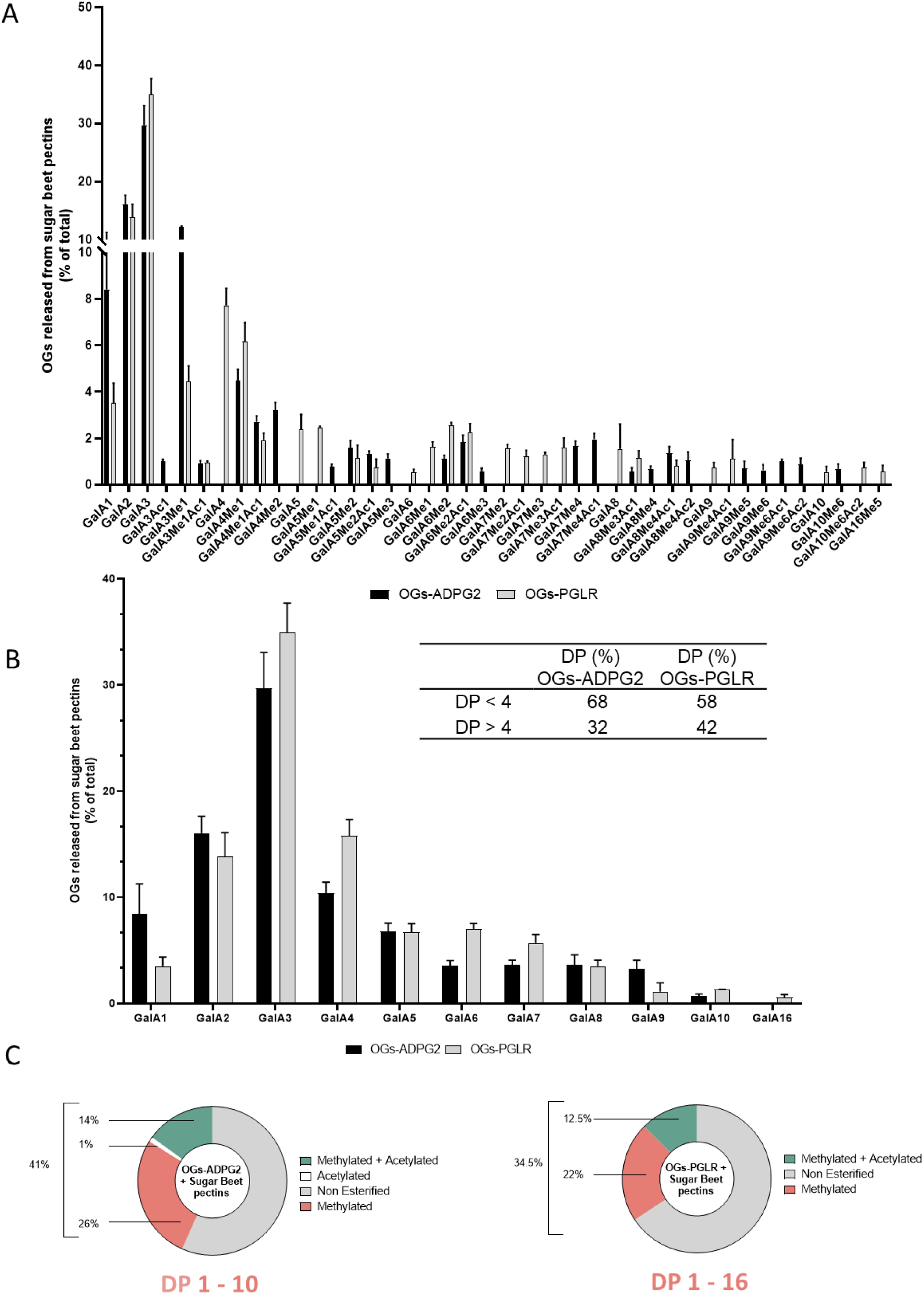
Characterization of oligogalacturonides (OGs) produced from sugar beet pectins (SBP): (A) OGs-ADPG2 (black) and OGs-PGLR (gray) comparisons. OGs with various DP, DM and DA are presented. (B) Determination of the proportion of OGs produced according to their DP. Inset: proportion of DP >4 or DP<4. (C) Determination of the proportion of esterified OGs

### 3.2. OGs-PGLR confer partial protection against wheat powdery mildew

Preventive wheat treatments with OGs-ADPG2 and OGs-PGLR pools were carried out 48 h before *Bgt* inoculation (**Fig. S1**). OGs were diluted at 5 g.L^-1^ in 10 mL of water, and the solution was sprayed on wheat leaves. Nine days after inoculation, disease symptoms were assessed by counting the number of *Bgt* colonies on the third leaf of each plant. OGs-PGLR application resulted in a reduction of the number of *Bgt* colonies on leaves, compared to the control (**Fig. 4A**). However, no significant reduction in the number of colonies was observed in plants treated with OGs-ADPG2 compared to the control (**Fig. 4B**). OGs-PGLR thus provided a significant protection rate of 31 % to wheat against powdery mildew disease (**Fig. 4B**).

**Fig. 4.**
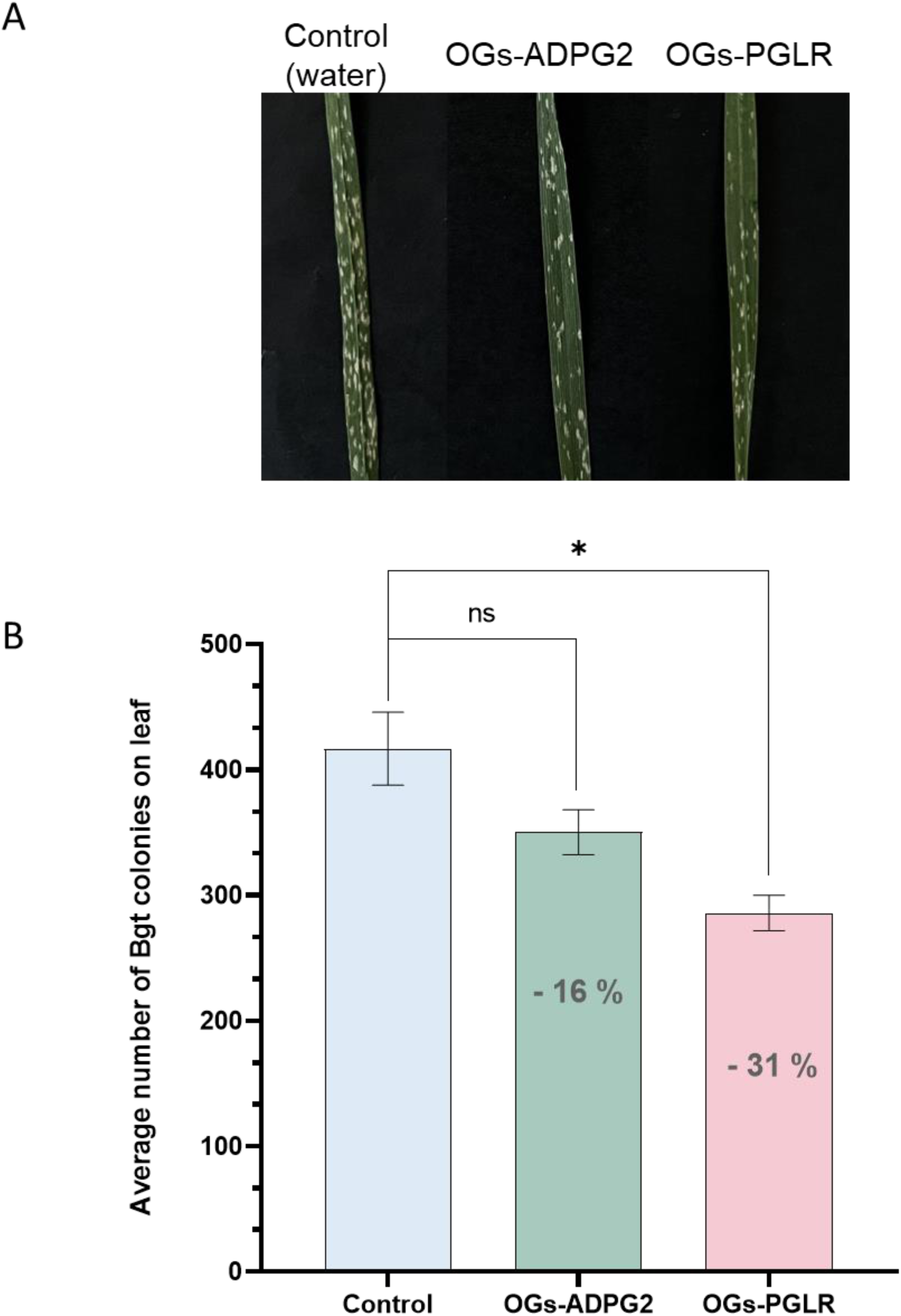
Average number of fungal colonies in response to OGs-ADPG2 and OGs-PGLR treatments, estimated on the third leaf of wheat plantlets cv. Alixan, 9 days after inoculation by *Bgt.* Plants were sprayed with 10 mL of OGs solution at 5 g.L^-1^ and inoculated by *Bgt* (5.10^5^ spores.mL^-1^) 2 days after treatments. (A) Symptoms (colonies) of *Bgt* on leaves according to the different treatments: Control (corresponding to water spraying), OGs-ADPG2 and OGs-PGLR. (B) Infection level of wheat leaves, after counting of *Bgt* colonies, according to the different treatments. The control condition corresponds to plants with water spraying as a treatment. T-test with Welch’s correction. *: P < 0.05 compared to the water control

### 3.3. OGs mechanism of action allowed partial protection against wheat powdery mildew

#### 3.3.1. OGs did not act directly on *Bgt* spore germination

To determine how OGs-PGLR protected wheat from powdery mildew, we first explored the potential bio-fungicide effect of these molecules. All molecules were assessed *in vitro* for their direct activity against *Bgt* spores. Plate assays revealed that none of the OG pools exhibited a direct antigerminative or antifungal effect against *Bgt* spores after 24 h of incubation, as no significant difference was observed in the various stages of spore development between treated and untreated conditions (**Fig. 5**). These findings are consistent with those reported by Randoux et al., (2010) for both acetylated and non-acetylated OGs, indicating that OGs, regardless of their DP, DM, or DA, do not exhibit any antigerminative effect. This suggests that the protection provided by OGs-PGLR is likely due to the induction of plant defense responses.

**Fig. 5.**
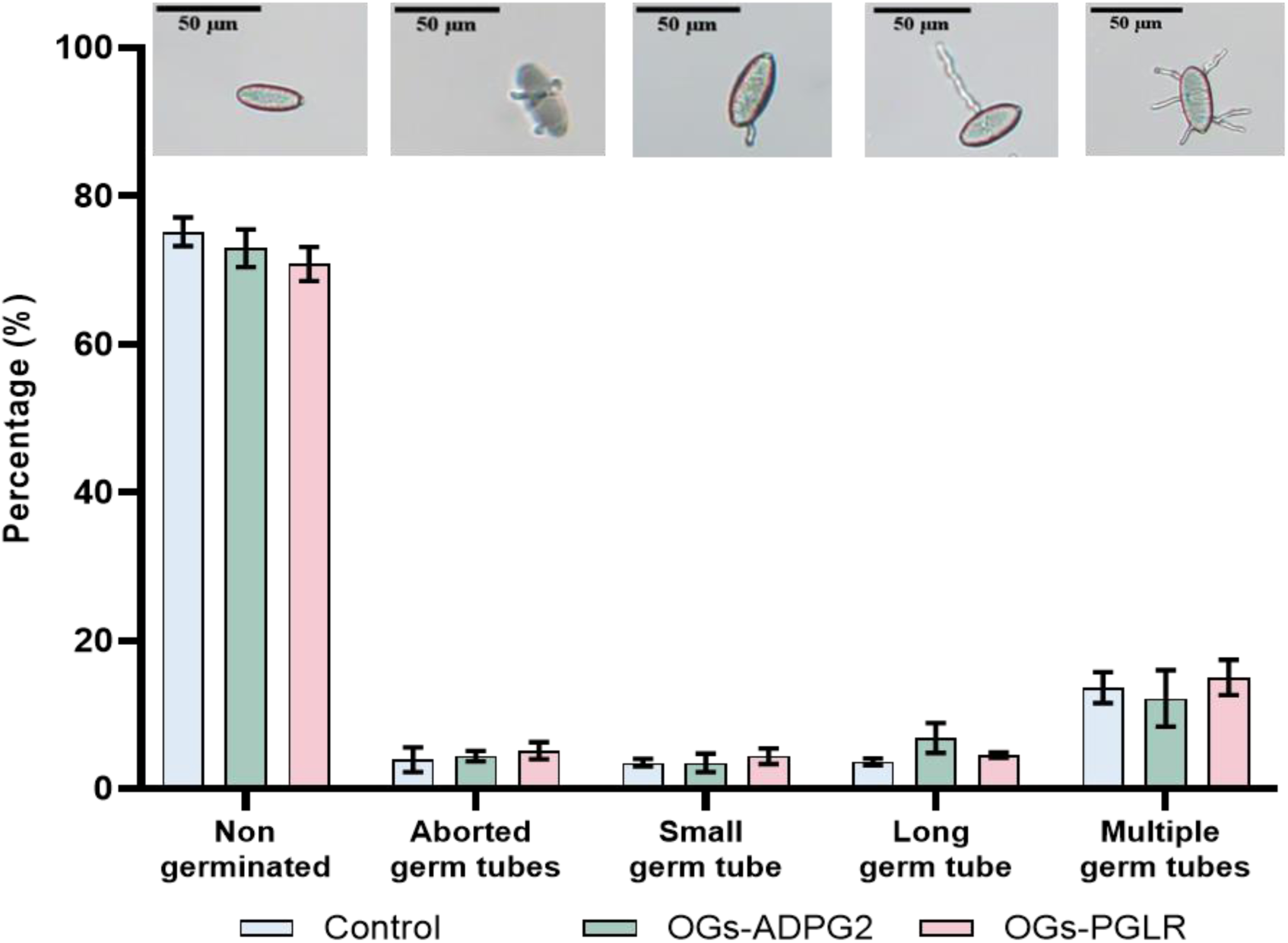
Effect of OGs-ADPG2 or OGs-PGLR on the germination of *Bgt* spores *in vitro.* Germination was assessed 24 h after the dispersion of the spores on Petri dishes containing agar (12 g.L^-1^) supplemented or not with OGs-ADPG2 et OGs-PGLR diluted at 5 g.L^-1^. The spores were divided into 5 categories: non germinated spores, spores with aborted germ tubes, one small germ tube, one long germ tube and spores with multiple germ tubes. Data represent means of 3 independent experiments. Statistical analysis with a multiple T-test was realized and showed no significant effect between the control plants and those treated with OGs-ADPG2 and OGs-PGLR.

#### 3.3.2. OGs induced defense-related mechanisms in wheat

The OGs produced, especially OGs-PGLR, offered partial protection to wheat against powdery mildew without directly affecting the pathogen’s spores. As a result, we shifted our focus to examining the indirect effects of these OGs on the plant to uncover the mechanisms triggered by OGs-PGLR that contribute to this partial protection, and to compare them with those induced by OGs-ADPG2.

The expression of targeted defense-related genes (**listed in Table S1**) was assessed using RT-qPCR at 48, 72, and 96 h post-treatment (hpt) with OGs (**Fig. S1**, **Fig. 6A**). Additionally, at 48 hpt with OGs, some plants were inoculated with *Bgt*. Gene expression was then evaluated 24 h (D1) and 48 h (D2) post-inoculation (**Fig. S1**, **Fig. 6B**). We determined a biological significance when the modification in expression data (log2FC) is greater than 1 or less than -1.

**Fig. 6.**
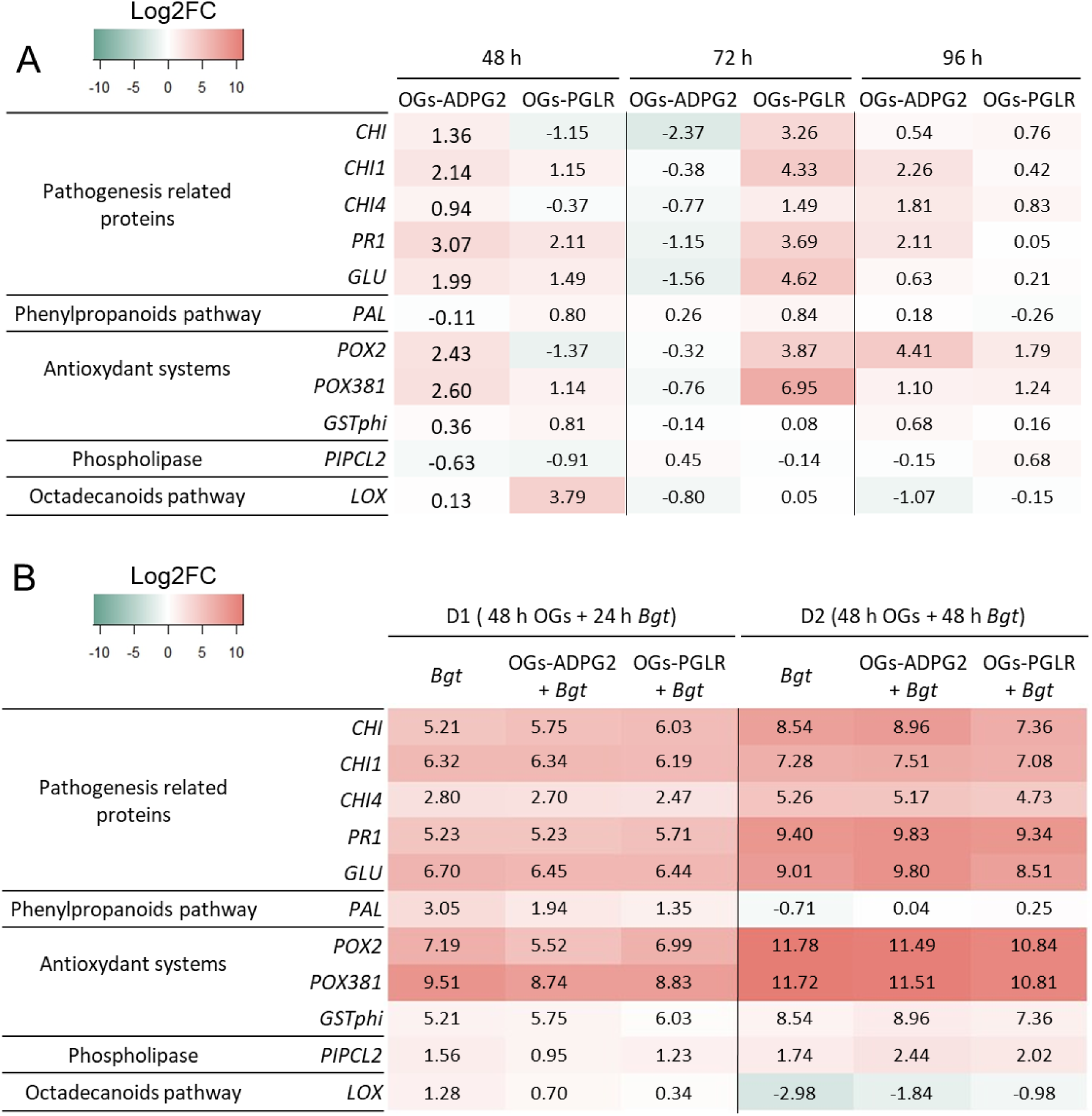
Pattern of defense gene expression, monitored at 48 h, 72 h and 96 h following the spraying of OGs-ADPG2, OGs-PGLR or water on the leaves of plants non-inoculated (A) or inoculated with *Bgt* 48h post treatment (B). Heatmap representation of gene expression fold changes, which is the mean of Log2 of 3 independent experiments. The results obtained were normalized by using TUB as the reference gene for each sample. Color scale represents the intensity of overexpression (pink) and down-regulation (green) of genes, as compared to control conditions (water-treatment and non-inoculated). Legends: *chitinase – CHI, basic chitinase 1 – CHI1, acidic chitinase (class 4) – CHI4, Pathogenesis related protein 1 – PR1, Glucanase – GLU, Phenyalanine ammonia lyase – PAL, Peroxidase 2 – POX2, Peroxidase 381 – POX381, Glutathion - S –Transferase (phi) – GSTphi, phosphoinositide specific Phospholipase – PIPCL2, Lipoxygenase– LOX*

Our results showed that both OGs pools triggered plant responses, mainly in non-inoculated conditions, but with different intensities at different time points (**Fig. 6A**).

In non-inoculated conditions and 48 hpt with OGs-ADPG2 (**Fig. 6A**), an overexpression of Pathogenesis related (PR) proteins (*CHI* – Log2FC = 1.36; *CHI1* – Log2FC = 2.14; *PR1* – Log2FC = 3.07; and *GLU* – Log2FC = 1.99) and peroxidases (*POX2* – Log2FC = 2.43; *POX381* – Log2FC = 2.60) was observed. For all other targeted-defense genes, expression levels remained close to that of control condition (**Fig. 6A**). At 48 hpt with OGs-PGLR (**Fig. 6A**), several PR protein genes were also overexpressed (*CHI1* – Log2FC = 1.15; *PR1* – Log2FC = 2.11; *GLU* – Log2FC = 1.49) along with a peroxidase (*POX381* – Log2FC = 1.14). In contrast to the OGs-ADPG2 treatment, OGs-PGLR induced a downregulation of *CHI* (Log2FC = -1.15) and *POX2* (Log2FC = -1.37) at 48 hpt (**Fig. 6A**). Finally, we observed that only OGs-PGLR treatment resulted in a significant overexpression of *LOX* (Log2FC = 3.79) at 48 hpt (**Fig. 6A**). At 72 hpt with OGs-ADPG2 (**Fig. 6A**), a downregulation of several target genes was observed, notably *CHI* (Log2FC = -2.37), *PR1* (Log2FC = -1.15), and *GLU* (Log2FC = -1.56), while the expression levels of other targeted defense genes returned close to control conditions. For OGs-PGLR at 72 hpt (**Fig. 6A**), the response differed, PR proteins (*CHI* – Log2FC = 3.26; *CHI1* – Log2FC = 4.33; *CHI4* – Log2FC = 1.49; *PR1* – Log2FC = 3.69; and *GLU* – Log2FC = 4.62) as well as peroxidases (*POX2* – Log2FC = 3.87; *POX381* – Log2FC = 6.95) were overexpressed. Expression levels at 48 hpt by OGs-PGLR for the other targeted genes remained close to control conditions (**Fig. 6A**). Finally, at 96 hpt with OGs-ADPG2 (**Fig. 6A**), an overexpression of *CHI1* (Log2FC = 2.26), *CHI4* (Log2FC = 1.81), *PR1* (Log2FC = 2.11), *POX2* (Log2FC = 4.41), and *POX381* (Log2FC = 1.10) was observed. In contrast for OGs-PGLR at 96 hpt (**Fig. 6A**), most targeted genes returned to control levels, except for the peroxidases *POX2* (Log2FC = 1.79) and *POX381* (Log2FC = 1.24) which remained overexpressed.

Under inoculated conditions (**Fig. 6B**), 24 hours after infection with *Bgt* (D1) (**Fig. 6B**, D1 and *Bgt* columns), all targeted defense genes were overexpressed, particularly the PR proteins (*CHI* – Log2FC = 5.21; *CHI1* – Log2FC = 6.32; *CHI4* – Log2FC = 2.80; *PR1* – Log2FC = 5.23; and *GLU* – Log2FC = 6.70) and peroxidases (*POX2* – Log2FC = 7.19; *POX381* – Log2FC = 9.51) (**Fig. 6B**, D1 and *Bgt* columns). We observed that at D1 post-inoculation, treatments with both OGs-ADPG2 and OGs-PGLR did not alter the expression of these genes compared to the *Bgt*-infected conditions (**Fig. 6B**). However, at D1, *PAL* expression decreased from a Log2FC of 3.05 with *Bgt* infection to a Log2FC of 1.94 in plants infected and treated with OGs-ADPG2 and to 1.35 in those infected and treated with OGs-PGLR (**Fig. 6B**, D1 column). Finally, *LOX* was overexpressed at D1 post-inoculation with *Bgt* (Log2FC = 1.28) but returned to control levels in plants infected and treated with OGs-ADPG2 (Log2FC = 0.70) or OGs-PGLR (Log2FC = 0.34) (Fig. 6B, D1 column). At 48 hours post-infection with *Bgt* (D2) (**Fig. 6B**, D2 and *Bgt* columns), PR proteins (*CHI* – Log2FC = 8.54; *CHI1* – Log2FC = 7.28; *CHI4* – Log2FC = 5.26; *PR1* – Log2FC = 9.40; and *GLU* – Log2FC = 9.01) and peroxidases (*POX2* – Log2FC = 11.78; *POX381* – Log2FC = 11.72) showed even greater overexpression (**Fig. 6B**, D2 and *Bgt* columns). At D2, we once again observed that the OGs-ADPG2 and OGs-PGLR treatments did not alter the response induced by *Bgt* (Fig. 6B). Additionally, at D2, *LOX* expression was significantly downregulated (Log2FC = -2.98) (**Fig. 6B**, D2 and *Bgt* columns). However, in plants infected and treated with OGs-ADPG2, the downregulation was less pronounced (Log2FC = -1.84), while those infected and treated with OGs-PGLR maintained an expression level close to the control conditions (Log2FC = -0.98) (**Fig. 6B**).

## 4. Discussion

In this study, we demonstrated that OGs produced through a mild, chemical-free process from a bio-sourced SBP can provide partial protection of wheat against powdery mildew, reducing foliar symptoms by 31%.

Our method of OGs production, derived from biosourced SBP, involves enzymatic hydrolysis using PGs. In addition to valorizing a by-product, the enzymatic approach we used allows more controlled hydrolysis, resulting in OGs with specific sizes and structures, without the need for chemical processes. In our case, this method was chosen for its reproducibility and its ability to maintain mild reaction conditions, which help preserve the natural esterifications present in the pectin, an important advantage over other OG production methods (Jayani et al., 2005). The most common method for OG production involves thermal hydrolysis of pectins (Einhorn-Stoll et al., 2020), which, while being relatively simple to implement and leading to the production of large quantities of OGs, it lacks specificity and reproducibility. Additionally, thermal degradation negatively affects the natural esterifications in pectin. For this reason, some have turned to chemical hydrolysis, which can also be used to produce OGs from pectin. This involves the use of acids or alkalis to achieve more controlled depolymerization. In addition, acidic treatments have also been used to acetylate OGs, such as those used in the study of Randoux et al., (2010) and Selim et al., (2017).

The precise characterization, and comparison, of the produced OGs allowed us to discriminate the OGs from the OGs-PGLR pool that could putatively induce protection. Based on our results, we first observed that OGs-PGLR are different from that of OGs-ADPG2 as previously described (Safran et al., 2023). We found that only 12 OGs were common to both batches of OGs (GalA1, GalA2, GalA3, GalA3Me1, GalA3Me1Ac1, GalA4Me1, GalA4Me1Ac1, GalA5Me2, GalA5Me2Ac1, GalA6Me2, GalA6Me2Ac1, GalA8Me3Ac1, GalA8Me4Ac1) (**Table S2**). Among the remaining OGs, 15 were specific to OGs-ADPG2, and the other 16 were specific to OGs-PGLR. These OGs specifically produced by PGLR (GalA4, GalA5, GalA5Me1, GalA6, GalA6Me1, GalA7Me2, GalA7Me2Ac1, GalA7Me3, GalA7Me3Ac1, GalA8, GalA9, GalA9Me4Ac1, GalA10, GalA10Me6Ac2, and GalA16Me5) could explain the different protective capacities as compared to OGs-ADPG2 (**Table S2**). We also observed that OGs-PGLR generally had a higher DP as compared to OGs-ADPG2 (**Fig. 3B**). Therefore, we can hypothesize that the higher proportion of OGs with DP > 4 plays a role in the ability of OGs-PGLR to protect wheat against powdery mildew. By comparison Randoux et al., (2010) use, on the same pathosystem, OGs with a high degree of polymerization (DP 2-25, mainly DP 4-7) and obtain also a protection. However, these OGs were produced with thermal degradation and needed a chemical intervention to acetylate the pools at 30%. Another study used OGs (DP 3-18, mainly DP 7-12) de-esterified by thermal degradation, against *Fusarium graminearum* resulted in a reduction of symptoms on the grains (Bigini et al., 2024).

In parallel, as a means to understand the protective effect of OGs-PGLR, we analysed the expression of some defense-related gene expression in wheat. Since OGs-PGLR did not directly affect the germination of *Bgt* spores (**Fig. 5**) or the expression of genes during infection (**Fig. 6B**), we can hypothesize that OGs-PGLR play a role before the inoculation at 48 hpt.

In wheat, our results showed that the expression of *CHI1*, *PR1*, and *GLU*, which code for PR proteins, as well as *POX381* (coding for a peroxidase) and *LOX* (coding for a lipoxygenase), is increased in response to OGs-PGLR starting at 48 hpt. By 72 hpt with OGs-PGLR, all targeted PR proteins and peroxidases are overexpressed, before returning close to control levels at 96 hpt. Thus, the overexpression of PR proteins and peroxidases in response to OGs treatment highlights the role of OGs in eliciting plant defense mechanisms. Previous studies indicate that PR proteins are indeed key players in plant defense against pathogens (Allario et al., 2023; Bindschedler et al., 2006; Saboki et al., 2011).

*PR1* was extensively studied and serves as a marker for plant defense responses. Its expression is often upregulated in response to pathogen attack and is associated with antifungal activity (Van Loon et al., 2006). *PR1* is also recognized as a marker for systemic acquired resistance in plants like *A. thaliana* (Zhang and Li, 2019). More recently, *PR1* was also overexpressed under non-infected conditions in wheat after the application of ulvan, a heteropolysaccharide (composed mainly of rhamnose, xylose, glucose, uronic acid, and sulfate) extracted from the cell walls of green algae (Velho et al., 2022). In addition, chitinases, encoded by *CHI* and *CHI1* genes, degrade chitin of fungal cell walls, which is essential for wheat’s defense against fungal infections (Grover, 2012). A previous study relative to the use of OGs against fungus *Botrytis cinerea* in grapevines also demonstrated a reduction in the lesions caused by the pathogen and an overexpression of PR proteins such as chitinases (Aziz et al., 2004). The action of chitinases was additive to that of glucanases, particularly β-1,3-glucanases encoded by *GLU*, which can degrade glucans found in fungal cell walls (Balasubramanian et al., 2012).

Peroxidases are also crucial enzymes in mediating the oxidative burst during pathogen attack (Bindschedler et al., 2006). They constitute one of the first lines of defense established by plants and contribute to the reinforcement of the cell wall at the sites of *Bgt* penetration (Allario et al., 2023). The observed upregulation of *POX2* at 48 hpt by OGs-PGLR or POX2 and *POX381* at 72 hpt by OGs-PGLR, or in inoculated conditions (D1 and D2), aligns with the role of peroxydases role in early defense mechanisms. Increased expression of *POX2* was observed in wheat leaves infected with *Bgt* (Båga et al., 1995) as well as with *Z. tritici* from (Ors et al., 2018). The expression of *POX381* was also found to be highly induced in wheat leaves following infection with *B. graminis f.sp. hordei* (Rebmann et al., 1991). Another study showed that the expression of *POX381* in wheat could be induced by bacterial lipopeptides, including surfactin and mycosubtilin, thus demonstrating elicitor activity (Khong et al., 2013). Similarly, OGs induced partial protection against *F. graminearum* and the overexpression of peroxidases was also demonstrated (Bigini et al., 2024). The induction of peroxidase activity was also observed following the application of OGs on alfalfa roots (Camejo et al., 2012).

However, the changes in expression of peroxidases and PR proteins at 48 and 72 hpt cannot solely explain the partial protection conferred by OGs-PGLR, as the expression of these genes was also induced by the application of OGs-ADPG2. Our results showed that only plants treated with OGs-PGLR showed a strong overexpression of a gene encoding a *LOX*, *9-LOX*. Moreover, under inoculated conditions with *Bgt* at D2, we observe that the expression of *9-LOX* was downregulated in the plants, while the it was at control-level in plants infected and treated with OGs-PGLR. Thus, we can hypothesize that LOX plays an important role in protection of wheat against *Bgt*. Oxylipin synthesis is initiated by lipoxygenases, and 9-LOX specifically oxidizes polyunsaturated fatty acids (PUFAs), such as linoleic and linolenic acids, at the 9th carbon leading to the formation of the corresponding fatty acid hydroperoxides (Feussner and Wasternack, 2002). Moreover, studies showed that the production of free fatty acid hydroperoxides via the 9-LOX pathway in tobacco plays a critical role in hypersensitive cell death triggered by cryptogein, a purified protein from *Phytophthora cryptogea* (Rustérucci et al., 1999). Another study also demonstrated the role of *9-LOX* as a regulator of root development and defense responses through a pathway independent of (+)-7-iso-jasmonic acid, which may be involved in triggering cell wall modifications (Vellosillo et al., 2007). A more recent study showed that *9-LOX* derivatives activate cell wall-based defense responses, particularly upstream of the brassinosteroid pathway, leading to the accumulation of callose (Marcos et al., 2015). Furthermore, the involvement of 9-LOX signaling pathways in inducing cell wall-based defense in *A. thaliana* was also confirmed by results obtained with *Golovinomyces cichoracearum*, a biotrophic fungus like *Bgt* (Marcos et al., 2015). Previous studies in wheat have demonstrated that the infiltration of SA into wheat leaves simultaneously triggers an increase in *LOX* gene expression and enhances lipoxygenase activity (Tayeh et al., 2016). Moreover, other studies have highlighted the importance of lipoxygenase activity induced by elicitors and, especially OGs, to trigger protection against wheat powdery mildew (Randoux et al., 2010; Tayeh et al., 2014; Velho et al., 2022).

Therefore, based on our various obervations, the application of the OGs-PGLR mainly acted mainly directly on the plant defense mechanisms, and especially on *LOX* expression. An additional priming activity was observed as *LOX* expression was maintained at a control level for plant infected and treated by OGs-PGLR, in contrast to untreated plants. A highest average DP in OGs-PGLR and the overexpression of *LOX* induced as early as 48 hpt seem therefore crucial for enabling wheat protection against powdery mildew.

## 5. Conclusion

In this study, we produced a mix of OGs from pectins extracted from sugar beet by-products, that were esterified (DM 62% and DA 27%). Hydrolysis of the pectin by two plant polygalacturonases, ADPG2 and PGLR, preserved the natural esterification, resulting in two distinct OG mixtures: OGs-ADPG2 (DP 1–10, 41% esterification) and OGs-PGLR (DP 1–16, 34.5% esterification). Only OGs-PGLR, with a higher average DP, provided partial protection to wheat against powdery mildew, reducing symptoms by 31%. Nevertheless, both OGs pools activated directly plant defense mechanisms at 48 et 72h for OG-ADPG2 and OG-PGLR, without any direct antigerminative effect on the fungus. Their activity involved mainly an up-regulation of peroxidase and PR protein encoding genes in non-infectious conditions. Furthermore, OGs-PGLR induced lipoxygenase (LOX) gene expression, indicating that the octadecanoid pathway may play a role in resistance induction by OGs-PGLR. No additional priming effect was observed for both OGs pools, excepted for *LOX*, whose down-regulation is attenuated in plant pre-treated with OGs-PGLR, under infectious conditions. Further investigations of this pathway could provide deeper insights into the mechanisms behind the enhanced protection provided by these OGs. Our findings highlight the potential value of biosourced OGs varying in size and esterification, as effective resistance inducers. This approach promotes the use of sugar beet by-products, reducing dependence on chemical pesticide and lowering agricultural carbon footprint. Finally, it opens up possibilities for local production of plant resistance inducers, based on the availability of pectin-rich by-products.

## Highlights

- OGs were produced from sugar beet by-products by two *A. thaliana* polygalacturonases (PGLR and ADPG2)
- OGs-PGLR (DP 1-16) induced 31 % wheat protection against powdery mildew in controlled conditions
- OGs-PGLR induced a direct overexpression of PR proteins, peroxidases and LOX encoding genes in plants, without any additional priming effect under infected conditions

## Funding sources

This work was supported by: Conseil Regional Hauts-de-France, Université de Picardie Jules Verne, Université du Littoral Côte d’Opale, CPER BiHauts Eco de France, and Alliance A2U (Artois, ULCO, UPJV) through a PhD grant awarded to Camille Carton.

## CRediT authorship contribution statement

**Camille Carton:** Writing – original draft, Methodology, Investigation, Formal analysis, Data curation, Conceptualization, **Josip Safran :** Methodology, Investigation, Formal analysis, **Sangeetha Mohanaraj:** Methodology, Investigation, Formal analysis, **Romain Roulard:** Methodology, Investigation, Formal analysis, **Jean-Marc Domon:** Methodology, Investigation, Conceptualization, **Solène Bassard :** Methodology, Investigation, **Natacha Facon :** Methodology, Investigation, **Benoît Tisserant :** Methodology, Investigation, **Gaelle Mongelard :** Methodology, Investigation, Conceptualization, **Laurent Gutierrez :** Methodology, Investigation, Conceptualization, **Beatrice Randoux :** Writing – review & editing, Methodology, Investigation, **Maryline Magnin-Robert:** Writing – review & editing, Methodology, Investigation, **Jérôme Pelloux:** Writing – review & editing, Conceptualization, **Corinne Pau-Roblot:** Writing – review & editing, Funding acquisition, Conceptualization, **Anissa Lounès-Hadj Sahraoui:** Writing – review & editing, Funding acquisition, Conceptualization.

## Declaration of competing interest

The authors declare that they have no conflicts of interest associated with this work.

## Acknowledgment

The authors would like to acknowledge Serge Pilard from the “Plateforme Analytique de l‘Université de Picardie Jules Verne” (Amiens, France) to LC-MS/MS experiments.

## Notes

### Competing Interest Statement

The authors have declared no competing interest.

## References

1. Albersheim, P., Darvill, A.G., Hcneil, M., Valent, B.S., Sharp, J.K., Nothnagel, E.A., Davis, K.R., Yamazaki, N., Gollin, D.J., York, W.S., Dudman, W.F., Darvill, J.E., Dell, A., 1983. Oligosaccharins: Naturally occurring carbohydrates with biological regulatory functions. Structure and Function of Plant Genomes 293–312.

2. Allario, T., Fourquez, A., Magnin-Robert, M., Siah, A., Maia-Grondard, A., Gaucher, M., Brisset, M.-N., Hugueney, P., Reignault, P., Baltenweck, R., Randoux, B., 2023. Analysis of Defense-Related Gene Expression and Leaf Metabolome in Wheat During the Early Infection Stages of Blumeria graminis f. sp. tritici. Phytopathology 113, 1537–1547. 10.1094/PHYTO-10-22-0364-R

3. Aziz, A., Heyraud, A., Lambert, B., 2004. Oligogalacturonide signal transduction, induction of defense-related responses and protection of grapevine against Botrytis cinerea. Planta 218, 767–774. 10.1007/s00425-003-1153-x

4. Bacete, L., Mélida, H., Miedes, E., Molina, A., 2018. Plant cell wall-mediated immunity: cell wall changes trigger disease resistance responses. Plant Journal 93, 614–636. 10.1111/tpj.13807

5. Båga, M., Chibbar, R.N., Kartha, K.K., 1995. Molecular cloning and expression analysis of peroxidase genes from wheat. Plant Mol Biol 29, 647–662. 10.1007/BF00041156

6. Balasubramanian, V., Vashisht, D., Cletus, J., Sakthivel, N., 2012. Plant β-1,3-glucanases: Their biological functions and transgenic expression against phytopathogenic fungi. Biotechnol Lett. 10.1007/s10529-012-1012-6

7. Bartlett, D.W., Clough, J.M., Godwin, J.R., Hall, A.A., Hamer, M., Parr-Dobrzanski, B., 2002. The strobilurin fungicides. Pest Manag Sci. 10.1002/ps.520

8. Bédouet, L., Courtois, B., Courtois, J., 2003. Rapid quantification of O-acetyl and O-methyl residues in pectin extracts, Carbohydrate Research.

9. Benedetti, M., Mattei, B., Pontiggia, D., Salvi, G., Savatin, D.V., Ferrari, S., 2017. Methods of isolation and characterization of oligogalacturonide elicitors, Methods in Molecular Biology. 10.1007/978-1-4939-6859-6_3

10. Bigini, V., Sillo, F., Giulietti, S., Pontiggia, D., Giovannini, L., Balestrini, R., Savatin, D. V, 2024. Oligogalacturonide application increases resistance to Fusarium head blight in durum wheat. J Exp Bot. 10.1093/jxb/erae050

11. Bindschedler, L. V., Dewdney, J., Blee, K.A., Stone, J.M., Asai, T., Plotnikov, J., Denoux, C., Hayes, T., Gerrish, C., Davies, D.R., Ausubel, F.M., Bolwell, G.P., 2006. Peroxidase-dependent apoplastic oxidative burst in Arabidopsis required for pathogen resistance. Plant Journal 47, 851–863. 10.1111/j.1365-313X.2006.02837.x

12. Bradford, M.M., 1976. A rapid and sensitive method for the quantitation of microgram quantities of protein utilizing the principle of protein-dye binding. Anal Biochem 72, 248–254. 10.1016/0003-2697(76)90527-3

13. Camejo, D., Martí, M.C., Olmos, E., Torres, W., Sevilla, F., Jiménez, A., 2012. Oligogalacturonides stimulate antioxidant system in alfalfa roots. Biol Plant 56, 537–544. 10.1007/s10535-012-0107-1

14. Curtis, T., Halford, N.G., 2014. Food security: The challenge of increasing wheat yield and the importance of not compromising food safety. Annals of Applied Biology. 10.1111/aab.12108

15. Einhorn-Stoll, U., Kastner, H., Fatouros, A., Krähmer, A., Kroh, L.W., Drusch, S., 2020. Thermal degradation of citrus pectin in low-moisture environment – Investigation of backbone depolymerisation. Food Hydrocoll 107. 10.1016/j.foodhyd.2020.105937

16. Ferrari, S., Savatin, D. V., Sicilia, F., Gramegna, G., Cervone, F., De Lorenzo, G., 2013. Oligogalacturonides: Plant damage-associated molecular patterns and regulators of growth and development. Front Plant Sci 4, 1–9. 10.3389/fpls.2013.00049

17. Feussner, I., Wasternack, C., 2002. The lipoxygenase pathway. Annu Rev Plant Biol. 10.1146/annurev.arplant.53.100301.135248

18. Galletti, R., Denoux, C., Gambetta, S., Dewdney, J., Ausubel, F.M., De Lorenzo, G., Ferrari, S., 2008. The AtrbohD-mediated oxidative burst elicited by oligogalacturonides in Arabidopsis is dispensable for the activation of defense responses effective against Botrytis cinerea. Plant Physiol 148, 1695–1706. 10.1104/pp.108.127845

19. Gamir, J., Minchev, Z., Berrio, E., García, J.M., De Lorenzo, G., Pozo, M.J., 2021. Roots drive oligogalacturonide-induced systemic immunity in tomato. Plant Cell Environ 44, 275–289. 10.1111/pce.13917

20. Godet, F., Limpert, E., 1998. Recent evolution of multiple resistance of Blumeria (Erysiphe) graminis f. sp. tritici to selected DMI and morpholine fungicides in France. Pestic Sci 54, 244–252. 10.1002/(SICI)1096-9063(1998110)54:3<244::AID-PS818>3.0.CO;2-8

21. Gravino, M., Savatin, D.V., MacOne, A., De Lorenzo, G., 2015. Ethylene production in Botrytis cinerea- and oligogalacturonide-induced immunity requires calcium-dependent protein kinases. Plant Journal 84, 1073–1086. 10.1111/tpj.13057

22. Grover, A., 2012. Plant Chitinases: Genetic Diversity and Physiological Roles. CRC Crit Rev Plant Sci 31, 57–73. 10.1080/07352689.2011.616043

23. Hocq, L., Guinand, S., Habrylo, O., Voxeur, A., Tabi, W., Safran, J., Fournet, F., Domon, J.M., Mollet, J.C., Pilard, S., Pau-Roblot, C., Lehner, A., Pelloux, J., Lefebvre, V., 2020. The exogenous application of AtPGLR, an endo-polygalacturonase, triggers pollen tube burst and repair. Plant Journal 103, 617–633. 10.1111/tpj.14753

24. Jayani, R.S., Saxena, S., Gupta, R., 2005. Microbial pectinolytic enzymes: A review. Process Biochemistry. 10.1016/j.procbio.2005.03.026

25. Joanna, B., Michal, B., Piotr, D., Agnieszka, W., Dorota, K., Izabela, W., 2018. Sugar Beet Pulp as a Source of Valuable Biotechnological Products, in: Advances in Biotechnology for Food Industry: Volume 14. Elsevier, pp. 359–392. 10.1016/B978-0-12-811443-8.00013-X

26. Kang, Y., Zhou, M., Merry, A., Barry, K., 2020. Mechanisms of powdery mildew resistance of wheat – a review of molecular breeding. Plant Pathol 69, 601– 617. 10.1111/ppa.13166

27. Khong, N.G., Randoux, B., Tisserant, B., Deravel, J., Tayeh, C., Coutte, F., Bourdon, N., Jacques, P., Reignault, P., 2013. Nonribosomal lipopeptides from Bacillus subtilis: potential resistance inducers and/or biopesticides for wheat crop?

28. Kumar, R., Choudhary, K., Bochalya, R.S., Sharma, D., Chandel, S., 2024. Effect of different sowing dates on growth and yield of wheat varieties. International Journal of Research in Agronomy 7, 17–24. 10.33545/2618060X.2024.v7.i9Sa.1414

29. Lakehal, A., Chaabouni, S., Cavel, E., Le Hir, R., Ranjan, A., Raneshan, Z., Novák, O., Păcurar, D.I., Perrone, I., Jobert, F., Gutierrez, L., Bakò, L., Bellini, C., 2019. A Molecular Framework for the Control of Adventitious Rooting by TIR1/AFB2-Aux/IAA-Dependent Auxin Signaling in Arabidopsis. Mol Plant 12, 1499–1514. 10.1016/j.molp.2019.09.001

30. Lecourieux, D., Mazars, C., Pauly, N., Ranjeva, R., Pugin, A., 2002. Analysis and effects of cytosolic free calcium increases in response to elicitors in Nicotiana plumbaginifolia cells. Plant Cell 14, 2627–2641. 10.1105/tpc.005579

31. Lemaire, A., Duran Garzon, C., Perrin, A., Habrylo, O., Trezel, P., Bassard, S., Lefebvre, V., Van Wuytswinkel, O., Guillaume, A., Pau-Roblot, C., Pelloux, J., 2020. Three novel rhamnogalacturonan I-pectins degrading enzymes from Aspergillus aculeatinus: Biochemical characterization and application potential. Carbohydr Polym 248, 116752. 10.1016/j.carbpol.2020.116752

32. Marcos, R., Izquierdo, Y., Vellosillo, T., Kulasekaran, S., Cascón, T., Hamberg, M., Castresana, C., 2015. 9-Lipoxygenase-derived oxylipins activate brassinosteroid signaling to promote cell wall-based defense and limit pathogen infection. Plant Physiol 169, pp.00992.2015. 10.1104/pp.15.00992

33. Meyers, E., Arellano, C., Cowger, C., 2019. Sensitivity of the U.S. Blumeria graminis f. sp. tritici Population to Demethylation Inhibitor Fungicides. Plant Dis 103, 3108–3116. 10.1094/PDIS-04-19-0715-RE

34. Miller, G.L., 1959. Use of Dinitrosalicylic Acid Reagent for Determination of Reducing Sugar. Anal Chem 31, 426–428. 10.1021/ac60147a030

35. Mohnen, D., 2008. Pectin structure and biosynthesis. Curr Opin Plant Biol 11, 266–277. 10.1016/j.pbi.2008.03.006

36. Mustafa, G., Khong, N.G., Tisserant, B., Randoux, B., Fontaine, J., Magnin-Robert, M., Reignault, P., Sahraoui, A.L.H., 2017. Defence mechanisms associated with mycorrhiza-induced resistance in wheat against powdery milde. Functional Plant Biology 44, 443–454. 10.1071/FP16206

37. Mwadzingeni, L., Shimelis, H., Dube, E., Laing, M.D., Tsilo, T.J., 2016. Breeding wheat for drought tolerance: Progress and technologies. J Integr Agric. 10.1016/S2095-3119(15)61102-9

38. Navazio, L., Moscatiello, R., Bellincampi, D., Baldan, B., Meggio, F., Brini, M., Bowler, C., Mariani, P., 2002. The role of calcium in oligogalacturonide-activated signalling in soybean cells. Planta 215, 596–605. 10.1007/s00425-002-0776-7

39. Ochoa-Meza, L.C., Quintana-Obregón, E.A., Vargas-Arispuro, I., Falcón-Rodríguez, A.B., Aispuro-Hernández, E., Virgen-Ortiz, J.J., Martínez-Téllez, M.Á., 2021. Oligosaccharins as elicitors of defense responses in wheat. Polymers (Basel) 13, 1–17. 10.3390/polym13183105

40. Ors, M.E., Randoux, B., Selim, S., Siah, A., Couleaud, G., Maumené, C., Sahmer, K., Halama, P., Reignault, P., 2018. Cultivar-dependent partial resistance and associated defence mechanisms in wheat against Zymoseptoria tritici. Plant Pathol 67, 561–572. 10.1111/ppa.12760

41. Pontiggia, D., Benedetti, M., Costantini, S., De Lorenzo, G., Cervone, F., 2020. Dampening the DAMPs: How Plants Maintain the Homeostasis of Cell Wall Molecular Patterns and Avoid Hyper-Immunity. Front Plant Sci. 10.3389/fpls.2020.613259

42. Putra, N.R., Rizkiyah, D.N., Abdul Aziz, A.H., Che Yunus, M.A., Veza, I., Harny, I., Tirta, A., 2023. Waste to Wealth of Apple Pomace Valorization by Past and Current Extraction Processes: A Review. Sustainability 15, 830. 10.3390/su15010830

43. Ralet, M.C., Cabrera, J.C., Bonnin, E., Quéméner, B., Hellìn, P., Thibault, J.F., 2005. Mapping sugar beet pectin acetylation pattern. Phytochemistry 66, 1832–1843. 10.1016/j.phytochem.2005.06.003

44. Randoux, B., Renard-Merlier, D., Mulard, G., Rossard, S., Duyme, F., Sanssené, J., Courtois, J., Durand, R., Reignault, P., 2010. Distinct defenses induced in wheat against powdery mildew by acetylated and nonacetylated oligogalacturonides. Phytopathology 100, 1352–1363. 10.1094/PHYTO-03-10-0086

45. Rasul, S., Dubreuil-Maurizi, C., Lamotte, O., Koen, E., Poinssot, B., Alcaraz, G., Wendehenne, D., Jeandroz, S., 2012. Nitric oxide production mediates oligogalacturonide-triggered immunity and resistance to Botrytis cinerea in Arabidopsis thaliana. Plant Cell Environ 35, 1483–1499. 10.1111/j.1365-3040.2012.02505.x

46. Rebmann, G., Hertig, C., Bull, J., Mauch, F., Dudler, R., 1991. Cloning and sequencing of cDNAs encoding a pathogen-induced putative peroxidase of wheat (Triticum aestivum L.), Plant Molecular Biology.

47. Rustérucci, C., Montillet, J.-L., Agnel, J.-P., Battesti, C., Alonso, B., Knoll, A., Bessoule, J.-J., Etienne, P., Suty, L., Blein, J.-P., Triantaphylidès, C., 1999. Involvement of Lipoxygenase-dependent Production of Fatty Acid Hydroperoxides in the Development of the Hypersensitive Cell Death induced by Cryptogein on Tobacco Leaves. Journal of Biological Chemistry 274, 36446–36455. 10.1074/jbc.274.51.36446

48. Saboki, E., Usha, K., Singh, B., 2011. Pathogenesis Related (PR) Proteins in Plant Defense Mechanism Age-Related Pathogen Resistance. Curr. Res. Technol. Adv. 1043–1054.

49. Safran, J., Tabi, W., Ung, V., Lemaire, A., Habrylo, O., Bouckaert, J., Rouffle, M., Voxeur, A., Pongrac, P., Bassard, S., Molinié, R., Fontaine, J., Pilard, S., Pau-roblot, C., Bonnin, E., Larsen, D.S., Morel-rouhier, M., Girardet, J., Lefebvre, V., Sénéchal, F., Mercadante, D., Pelloux, J., 2023. Plant polygalacturonase structures specify enzyme dynamics and processivities to fine-tune cell wall pectins. Plant Cell 1–19. 10.1093/plcell/koad134

50. Sakamoto, T., Sakai, T., 1995. Analysis of structure of sugar-beet pectin by enzymatic methods. Phytochemistry 39, 821–823. 10.1016/0031-9422(95)00979-H

51. Selim, S., Sanssené, J., Rossard, S., Courtois, J., 2017. Systemic induction of the defensin and phytoalexin pisatin pathways in pea (Pisum sativum) against aphanomyces euteiches by acetylated and nonacetylated oligogalacturonides. Molecules 22, 1–17. 10.3390/molecules22061017

52. Shahin, L., Zhang, L., Mohnen, D., Urbanowicz, B.R., 2023. Insights into pectin O-acetylation in the plant cell wall: structure, synthesis, and modification. The Cell Surface 9, 100099. 10.1016/j.tcsw.2023.100099

53. Silva-Sanzana, C., Estevez, J.M., Blanco-Herrera, F., 2020. Influence of cell wall polymers and their modifying enzymes during plant–aphid interactions. J Exp Bot 71, 3854–3864. 10.1093/jxb/erz550

54. Spinei, M., Oroian, M., 2023. Structural, functional and physicochemical properties of pectin from grape pomace as affected by different extraction techniques. Int J Biol Macromol 224, 739–753. 10.1016/j.ijbiomac.2022.10.162

55. Spinei, M., Oroian, M., 2022. The Influence of Extraction Conditions on the Yield and Physico-Chemical Parameters of Pectin from Grape Pomace. Polymers (Basel) 14. 10.3390/polym14071378

56. Tayeh, C., Randoux, B., Laruelle, F., Bourdon, N., Reignault, P., 2016. Phosphatidic acid synthesis, octadecanoic pathway and fatty acids content as lipid markers of exogeneous salicylic acid-induced elicitation in wheat. Functional Plant Biology 43, 512–522. 10.1071/FP15347

57. Tayeh, C., Randoux, B., Vincent, D., Bourdon, N., Reignault, P., 2014. Exogenous trehalose induces defenses in wheat before and during a biotic stress caused by powdery mildew. Phytopathology 104, 293–305. 10.1094/PHYTO-07-13-0191-R

58. Thakur, M., Sohal, B.S., 2013. Role of Elicitors in Inducing Resistance in Plants against Pathogen Infection: A Review. ISRN Biochem 2013, 1–10. 10.1155/2013/762412

59. Usmani, Z., Sharma, M., Diwan, D., Tripathi, M., Whale, E., Jayakody, L.N., Moreau, B., Thakur, V.K., Tuohy, M., Gupta, V.K., 2022. Valorization of sugar beet pulp to value-added products: A review. Bioresour Technol. 10.1016/j.biortech.2021.126580

60. Van Loon, L.C., Rep, M., Pieterse, C.M.J., 2006. Significance of inducible defense-related proteins in infected plants. Annu Rev Phytopathol. 10.1146/annurev.phyto.44.070505.143425

61. Vandesompele, J., De Preter, K., Pattyn, F., Poppe, B., Van Roy, N., De Paepe, A., Speleman, F., 2002. Accurate normalization of real-time quantitative RT-PCR data by geometric averaging of multiple internal control genes. Genome Biol 3, research0034.1. 10.1186/gb-2002-3-7-research0034

62. Velho, A.C., Dall’Asta, P., de Borba, M.C., Magnin-Robert, M., Reignault, P., Siah, A., Stadnik, M.J., Randoux, B., 2022. Defense responses induced by ulvan in wheat against powdery mildew caused by Blumeria graminis f. sp. tritici. Plant Physiology and Biochemistry 184, 14–25. 10.1016/j.plaphy.2022.05.012

63. Vellosillo, T., Martínez, M., López, M.A., Vicente, J., Cascón, T., Dolan, L., Hamberg, M., Castresana, C., 2007. Oxylipins produced by the 9-lipoxygenase pathway in Arabidopsis regulate lateral root development and defense responses through a specific signaling cascade. Plant Cell 19, 831– 846. 10.1105/tpc.106.046052

64. Vielba-Fernández, A., Polonio, Á., Ruiz-Jiménez, L., de Vicente, A., Pérez-García, A., Fernández-Ortuño, D., 2020. Fungicide resistance in powdery mildew fungi. Microorganisms. 10.3390/microorganisms8091431

65. Voxeur, A., Habrylo, O., Guénin, S., Miart, F., Soulié, M.C., Rihouey, C., Pau-Roblot, C., Domon, J.M., Gutierrez, L., Pelloux, J., Mouille, G., Fagard, M., Höfte, H., Vernhettes, S., 2019. Oligogalacturonide production upon Arabidopsis thaliana-Botrytis cinerea interaction. Proc Natl Acad Sci U S A 116, 19743–19752. 10.1073/pnas.1900317116

66. Willats, W.G.T., Knox, J.P., Mikkelsen, J.D., 2006. Pectin: New insights into an old polymer are starting to gel. Trends Food Sci Technol 17, 97–104. 10.1016/j.tifs.2005.10.008

67. Winning, H., Viereck, N., Nørgaard, L., Larsen, J., Engelsen, S.B., 2007. Quantification of the degree of blockiness in pectins using 1H NMR spectroscopy and chemometrics. Food Hydrocoll 21, 256–266. 10.1016/j.foodhyd.2006.03.017

68. Zeuner, B., Thomsen, T.B., Stringer, M.A., Krogh, K.B.R.M., Meyer, A.S., Holck, J., 2020. Comparative Characterization of Aspergillus Pectin Lyases by Discriminative Substrate Degradation Profiling. Front Bioeng Biotechnol 8. 10.3389/fbioe.2020.00873

69. Zhang, Y., Li, X., 2019. Salicylic acid: biosynthesis, perception, and contributions to plant immunity. Curr Opin Plant Biol. 10.1016/j.pbi.2019.02.004

70. Zhang, Y., Yu, J., Wang, X., Durachko, D.M., Zhang, S., Cosgrove, D.J., 2021. Molecular insights into the complex mechanics of plant epidermal cell walls. Science (1979) 372, 706–711. 10.1126/science.abf2824

71. Zhao, C., Wu, C., Li, K., Kennedy, J.F., Wisniewski, M., Gao, L., Han, C., Liu, J., Yin, H., Wu, X., 2022. Effect of Oligogalacturonides on Seed Germination and Disease Resistance of Sugar Beet Seedling and Root. Journal of Fungi 8, 716. 10.3390/jof8070716

72. Zou, S., Xu, Y., Li, Q., Wei, Y., Zhang, Y., Tang, D., 2023. Wheat powdery mildew resistance: from gene identification to immunity deployment. Front Plant Sci 14, 1–7. 10.3389/fpls.2023.1269498

