## Supplemental Table 1 for "Oligogalacturonides, produced by enzymatic degradation of sugar beet by-products, confer partial protection against wheat powdery mildew"

**Table S1.** Primers of genes analyzed by real-time reverse-transcription polymerase chain reaction (RT-qPCR)

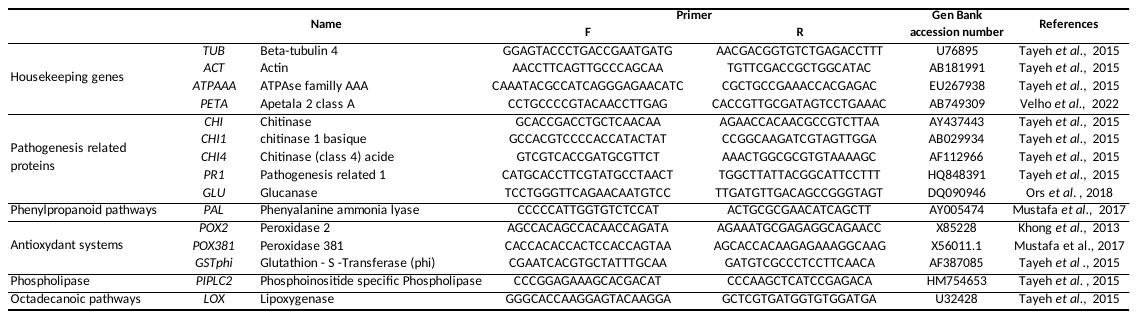
