## Supplemental Table 2 for "Oligogalacturonides, produced by enzymatic degradation of sugar beet by-products, confer partial protection against wheat powdery mildew"

**Table S2.** OGs panel produced specifically by ADPG2 and PGLR. Legend: + corresponding to present in the pools.

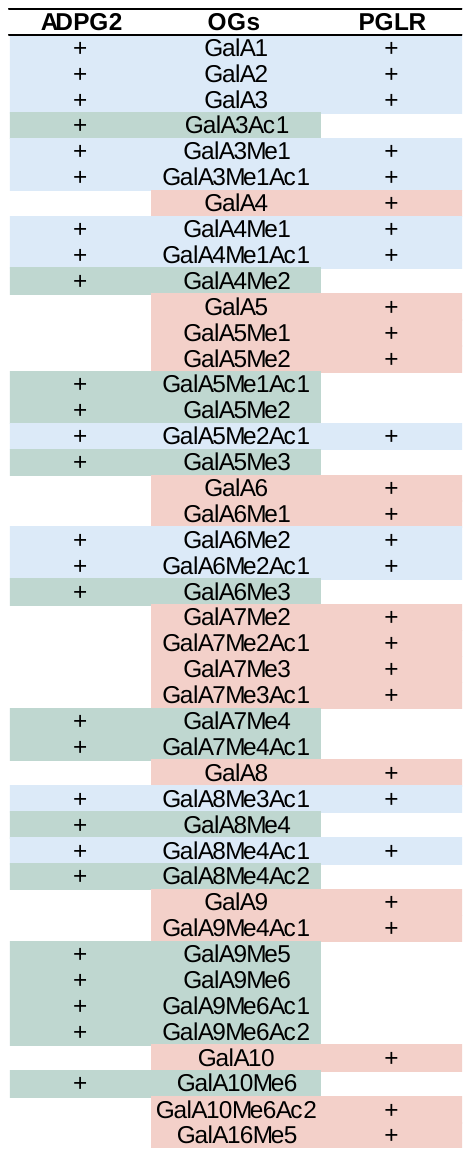
