## Supplemental Fig S1 for "Oligogalacturonides, produced by enzymatic degradation of sugar beet by-products, confer partial protection against wheat powdery mildew"

**Fig. S1**. Procedure for the evaluation test of wheat protection against powdery mildew following preventive application with OGs. Wheat leaf samples were harvested 48 h post-treatment (hpt) with OGs or dH_2_O (before *Bgt* inoculation), as well as 24 h (D1) and 48 h (D2) post-inoculation with *Bgt* (equivalent to 72 and 96 hpt under non-infectious conditions).

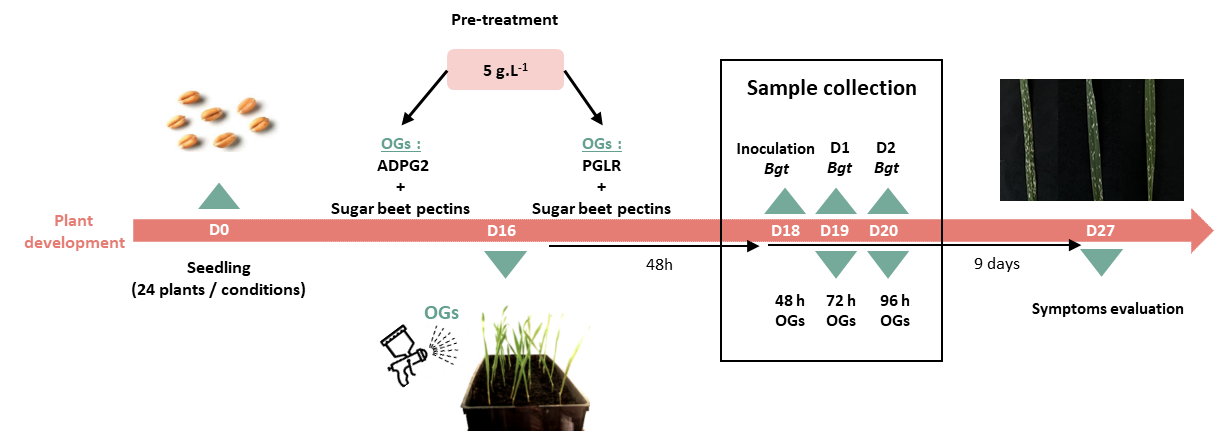
